# Differences in Brain Microtubule Electrical Activity of the Hippocampus and Neocortex from the Adult Rat

**DOI:** 10.1101/2025.11.18.688943

**Authors:** Horacio F. Cantiello, Cintia Y. Porcari, Virginia H. Albarracín, David Murphy, Andre S. Mecawi, Andrea Godino, María del Rocío Cantero

## Abstract

Microtubules (MTs) are essential cytoskeletal structures in neurons that generate electrical oscillations in the frequency range of mammalian brain waves. However, the role of these MT oscillations in brain function remains largely unknown. Here, we sought to gain insight into MT electrical oscillatory activity from different brain regions with specific functions, the hippocampus and the neocortex from the adult rat brain. We obtained local field potentials (LFP) from the frozen brain regions under non-depolarized (high external NaCl) and depolarized (high external KCl) saline solutions, observing spontaneous oscillations under both conditions. The electrical oscillations of the brain tissue had different amplitudes in the absence (0 mV) or presence (100 mV) of holding potential and were inhibited by the MT stabilizer paclitaxel. A frequency domain spectral analysis of the time records revealed the presence of two major peaks at approximately ∼38 Hz and ∼93 Hz in both preparations. However, the energy contribution of each peak was different in the hippocampus compared to neocortex. Coupled with our electron microscopy observations, these data suggest that rat brain MTs produce electrical oscillations with specific properties in the various regions of the mammalian brain, which could be partially related as their intra-axonal distributions. MT oscillations may be implicated in the wave coherence of brain activity, supporting their contribution to the concept of a brain central oscillator that drives its function.

## Introduction

Cellular electrical activity involves resting and action potentials associated with ionic movements across the cell membrane, whose function is central for the activity of such organs as the brain. However, contributions made by intracellular elements such as the cytoskeleton in the electrical activity of neurons are not understood.

Brain waves are thought to reflect synchronized electrical oscillations generated by the activity of large ensembles of active neurons, which can be observable through electroencephalogram (EEG) and local field potential (LFP) recordings^1-3^. Brain waves are found across various animal phyla, suggesting a fundamental role in brain computations. Different types of brain waves, characterized by frequency and amplitude such as alpha, beta, delta, theta, and gamma waves, exhibit diverse patterns of oscillatory activity influenced by tasks, cognitive states, and other factors. Coherence among different brain regions, indicating synchronization or coordination of electrical activity, is crucial in cognitive functions including attention, memory, and perception^4-10^. For example, brain wave coherence between frontal and parietal regions in the alpha and beta frequency ranges is associated with improved attentional performance and working memory tasks^11-12^. Wave synchronization depends on brain regions with specific wave form frequency and amplitude linked to sensory and physiological responses^9,13,14^. Intracerebral recordings have unveiled synchronization of high-frequency oscillations across widely distributed brain regions^15,16^.

Recent studies highlight the importance of wave coherence in various cognitive functions, particularly attention and memory^10^. Increased coherence between specific brain regions is linked to improved attentional performance, while decreased coherence is associated with attention deficits. The coupling of patterns between hippocampus and cortical activity, for example, facilitates memory consolidation processes. Short-term memory maintenance involves coordinated activity between the prefrontal cortex and hippocampus, with distinct neuronal populations encoding different aspects of memory^17^. Although brain wave synchronization may recruit synaptic function and neuronal channel activity^18-21^ brain wave function involves travelling waves^22^ that transcend neuronal function.

Previous studies have revealed that cytoskeletal polymers, including actin filaments and microtubules (MTs) have intrinsic electrical properties that contribute to intracellular circuits in neurons^23,24^ and possibly electrical interfaces with ion channels implicated in neuronal activity^25-27^. In particular, brain MTs generate spontaneous electrical oscillations exhibiting frequencies similar to brain waves^28^, suggesting a central role in brain function^23,24,29^ that underscore the capacity to generate oscillations observed across various mammalian species. The observation of MT-originated oscillations in permeabilized murine hippocampal neurons and the ex vivo honeybee brain with frequencies within the typical brain wave range led us to postulate the existence of a brain central oscillator based on elements of the cytoskeleton^23,24^. Interestingly, super-resolution microscopy of osmosensory neurons isolated from the supraoptic nuclei of adult rats showed diverse complexity of MT networks^30^, potentially influencing resting oscillatory waves and coherence in the brain.

Here we used LFPs with a current-to-voltage patch clamp system to explore the presence of endogenous oscillations in identified regions of the adult rat brain, revealing specific patterns with distinct frequency peaks that resemble other brain MT preparations^23,29^. Wave form analysis also suggested a partial similarity to oscillations observed in isolated mammalian brain MTs, implying a contribution of the cytoskeleton in mediating intracellular electrical signals. Understanding the mechanisms underlying these phenomena is crucial for deeper insights into brain function and dysfunction. In a companion manuscript, we also provide evidence that specific brain domain frequency peaks may contribute to a physiological response under non-homeostatic salt conditions^31^.

## Results

### Electrical activity of adult rat brain tissue

To obtain electrical information from the rat brain, tissue matter was obtained, processed, and kept frozen at ࢤ20°C until the experiment, where 1-2 mm size samples were defrosted into normal (high NaCl) saline solution. We conducted further experiments in a Ca^2+^-free “intracellular-like” solution containing high KCl (140 mM) and 1 mM EGTA.

### Electrical activity of rat brain tissue in the presence of KCl in the bath solution

Rat brain tissue samples (n = 43) maintained in extracellular solution until the moment of the experiment were tested with a patch clamp amplifier (Fig. 1) and a pipette electrode filled with an intracellular type of saline, as previously reported^24^, as described in Materials and Methods. In this study, the voltage-clamp patch pipette was used to assess local field potentials (LFP) in the form of electric currents at the location of the pipette under symmetrical ionic conditions. The tip conductance was first measured in saline solution (Fig 2A), which was highly linear as expected (data not show) and devoid of any oscillatory behavior, except for the line frequency at ∼50 Hz (Fig 2A, Right). The tip resistance of the patch pipette was 5.76 ± 1.38 MΩ (n = 32) under symmetrical saline (identical KCl concentration in pipette and bath) conditions.

**Figure 1.**
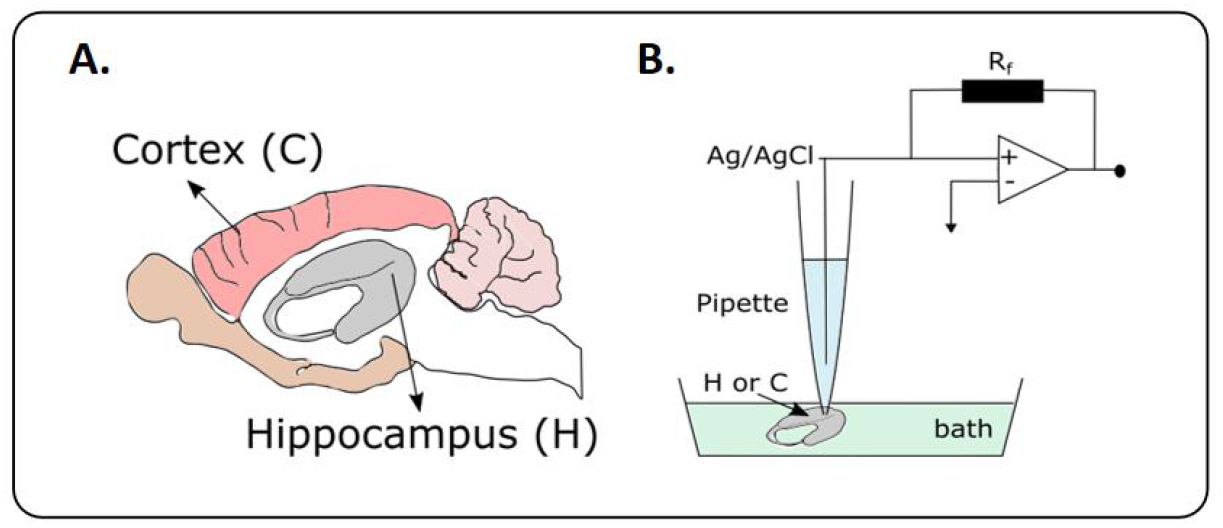
Experimental setup. A. Brain schematic indicating the cortex and the hippocampus localization. B. Technical outline showing the loose-patch data acquisition system.

**Figure 2.**
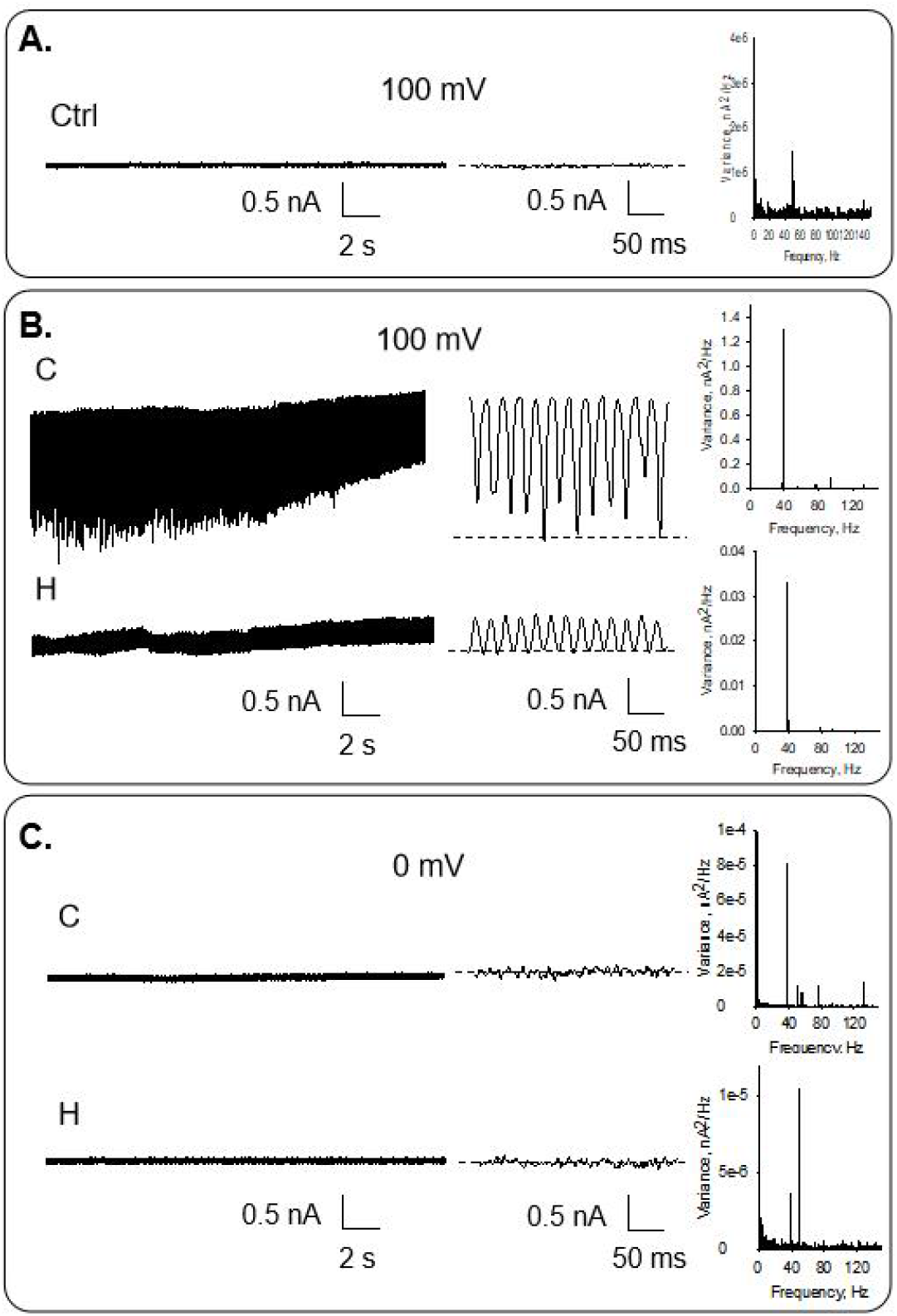
Electrical activity of rat brain tissue incubated in extracellular solution. A. Electrical recording of a free pipette in KCl bath solution at 100 mV as a negative control. B. Electrical recording of cortex (C) and hippocampus (H) with a KCl bathing solution. Data were obtained at 100 mV. C. Electrical recording of C and H with a KCl bathing solution obtained at zero mV. Experiments are representative of 7 and 5 for C and H, respectively. *(Left*. Unfiltered tracing. *Medium*. Expansion of a region on the left. *Right*. Power spectra).

Voltage-clamped cortex samples displayed spontaneous, self-sustained electrical oscillations that responded directly to the magnitude and polarity of the electrical stimulus (Fig 2B Top). Similar findings were obtained with the hippocampus samples (Fig. 2B Bottom). Fourier spectra showed a fundamental frequency of ∼38 Hz. for both samples (cortex and hippocampus, Fig. 2B Right). The frequency is present even at 0 mV, indicating the presence of a chemical gradient between the surrounding media of the brain matter and the inside of the pipette patch (Fig. 2C). The oscillation was more prominent for the cortex samples than for the hippocampus. Three-dimensional phase-space portraits showed limit cycles, evidencing mono-periodic oscillatory behavior at 100 mV (Fig. 4A) for both hippocampus and cortex.

### Electrical activity of rat brain tissue after incubation in KCl

To evaluate the effect of ionic composition on the electrical currents, rat brain tissue was incubated in KCl solution (see Methods) for 24-48 h. The tip conductance was first measured in the bath solution as a control (Fig 3A) and devoid of any oscillatory behavior, except for the line frequency at ∼50 Hz (Fig 3A, Right). As before, the brain matter was tested with a patch-clamp amplifier under symmetrical KCl conditions. Voltage-clamped cortex samples displayed spontaneous, self-sustained electrical oscillations that responded directly to the magnitude and polarity of the electrical stimulus (Fig 4b Top). Similar findings were obtained with the hippocampus samples (Fig. 3B). Fourier spectra showed a fundamental frequency of ∼38 Hz for both samples (cortex and hippocampus, Fig. 3B Right). The oscillatory frequency is present even at 0 mV, indicating that the presence of spontaneous fluctuations in the chemical gradient between the surrounding media of the brain matter and the inside of the pipette patch (Fig. 4C) are sufficient to elicit electrical oscillations as recently calculated for isolated brain MTs^32^.

**Figure 3.**
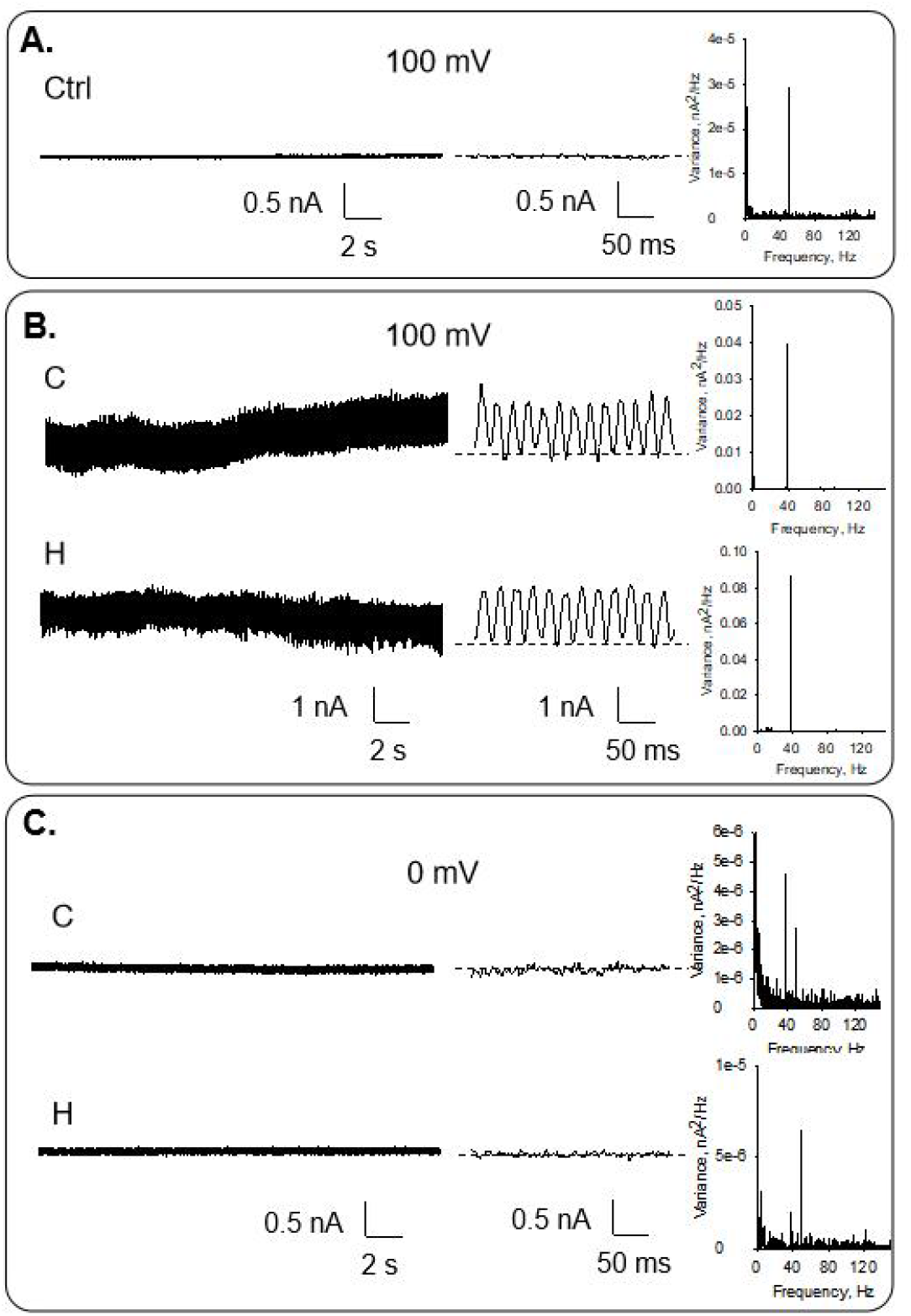
Electrical activity of rat brain tissue incubated in KCl solution. A. Electrical recording of a free pipette in KCl bath solution at 100 mV as a negative control. B. Electrical recording of cortex (C) and hippocampus (H) with a KCl bathing solution. Data were obtained at 100 mV. C. Electrical recording of C and H with a KCl bathing solution obtained at zero mV. Experiments are representative of 9 and 7 for C and H, respectively. (*Left*. Unfiltered tracing. *Medium*. Expansion of a region on the left. *Right*. Power spectra).

**Figure 4.**
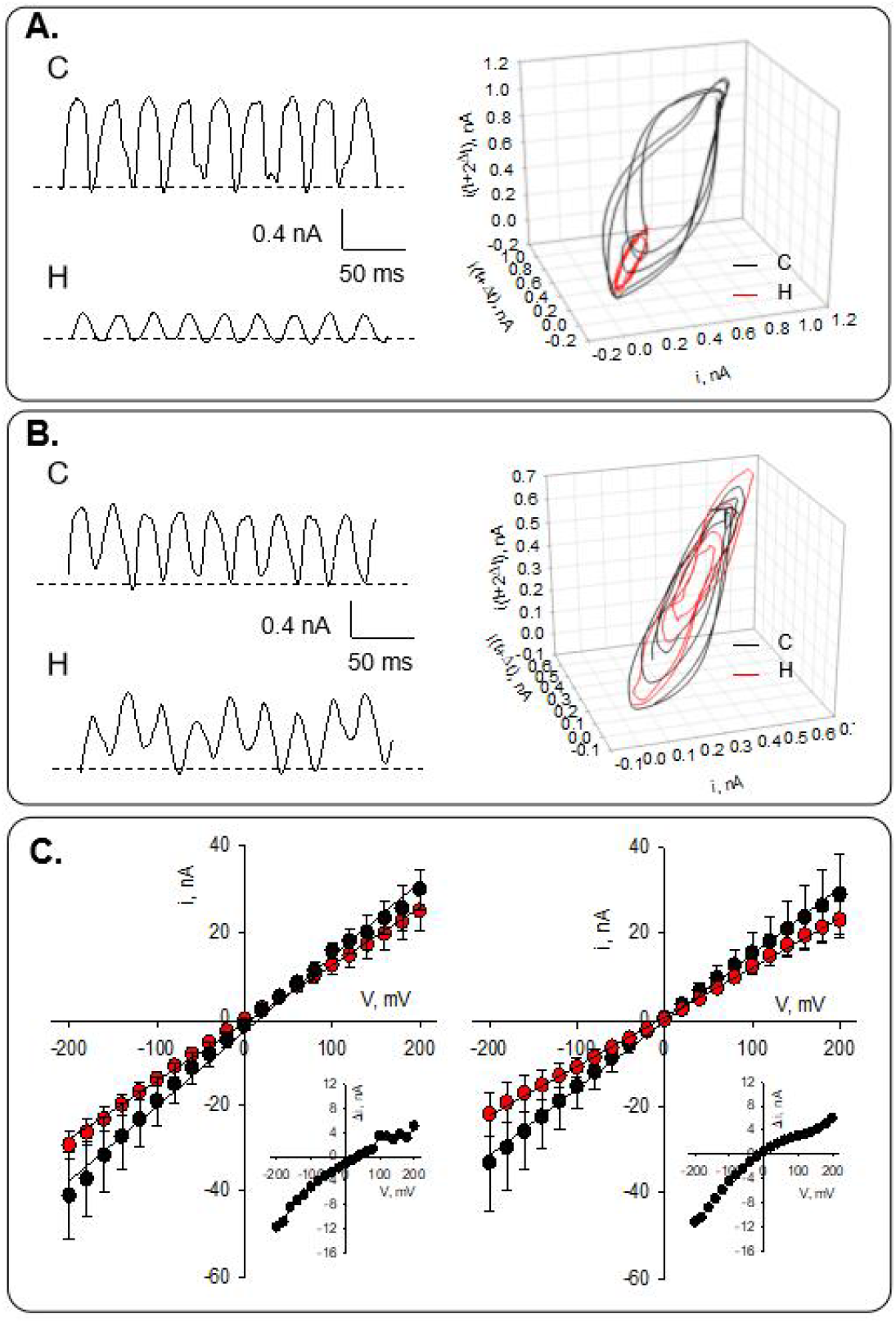
Current oscillations and current-to-voltage analysis. A. Left. Representative tracings at 100 mV compared the recordings obtained in cortex (C) and hippocampus (H) after incubating the samples in an extracellular solution and patching them in symmetrical KCl. Right. As indicated, three-dimensional phase-space portraits show monoperiodic limit cycles for the cortex and hippocampus. The delay time for the first and second derivatives adopted for phase portraits was 10 ms. B. Left. Representative tracings at 100 mV compared the recordings obtained in C and H after incubating the samples in KCl solution and patching in symmetrical KCl. Right. As indicated, Three-dimensional phase-space portraits show monoperiodic limit cycles for the cortex and hippocampus. The delay time for the first and second derivatives adopted for phase portraits was 10 ms. C. Current-to-voltage obtained for C (Left) and H(Right), showing in Black symbols the response after incubation with extracellular solution and in red symbols incubation with KCl (24-48 h). The inserts show the difference between Red-Black conditions, indicating similar inward rectification.

In contrast with the observation reported in the previous section, after incubation with KCl the oscillations were similar for the cortex and hippocampus samples (Fig. 4B). The three-dimensional phase-space portraits showed limit cycles, evidencing mono-periodic oscillatory behavior at 100 mV for both samples.

The current-to-voltage relationship was approximately linear when samples were incubated in the extracellular (High Na^+^) solution (Fig. 4C, Black symbols). Mean conductance under this condition was 173.3 ± 3.1 nS (n = 4) for cortex, and 154.8 ± 1.6 nS (n = 4) for hippocampus. The conductances were statistically different with p = 0.0018, compared by t-test. The conductance also differed after sample incubation with KCl (Fig. 4C, Red symbols vs. Black symbols). Under this condition, the mean conductance was 134.3 ± 1.4 nS (n = 3) for cortex, and 113.9 ± 0.8 nS (n = 6) for hippocampus. Again, the conductances were statistically different from each other (p < 0.0001), and also when compared with the incubation with extracellular solution (173.3 nS vs 134.3 nS p = 0.0002, and 154.8 nS vs 113.9 nS, p < 0.0001).

### Role of the microtubules in the electrical oscillations of brain matter

To evaluate the contribution of MTs to the rat brain electrical oscillations, we assessed the effect of the MT stabilizer paclitaxel that eliminates the electrical oscillations of MT preparations^23,29^. Paclitaxel (1.5 mM) was added after recording the electrical oscillations of the preparation in bathing KCl solution. The spontaneous electrical oscillations were significantly reduced after the addition of paclitaxel, as shown in Fig. 5 (n = 3), to reach an almost complete inhibition in the case of the hippocampus. Interestingly, the frequency of 38 Hz is preserved at its minimum after paclitaxel addition. These results imply that MTs are fundamental participants of the intracellular oscillations observed in the rat brain.

**Figure 5.**
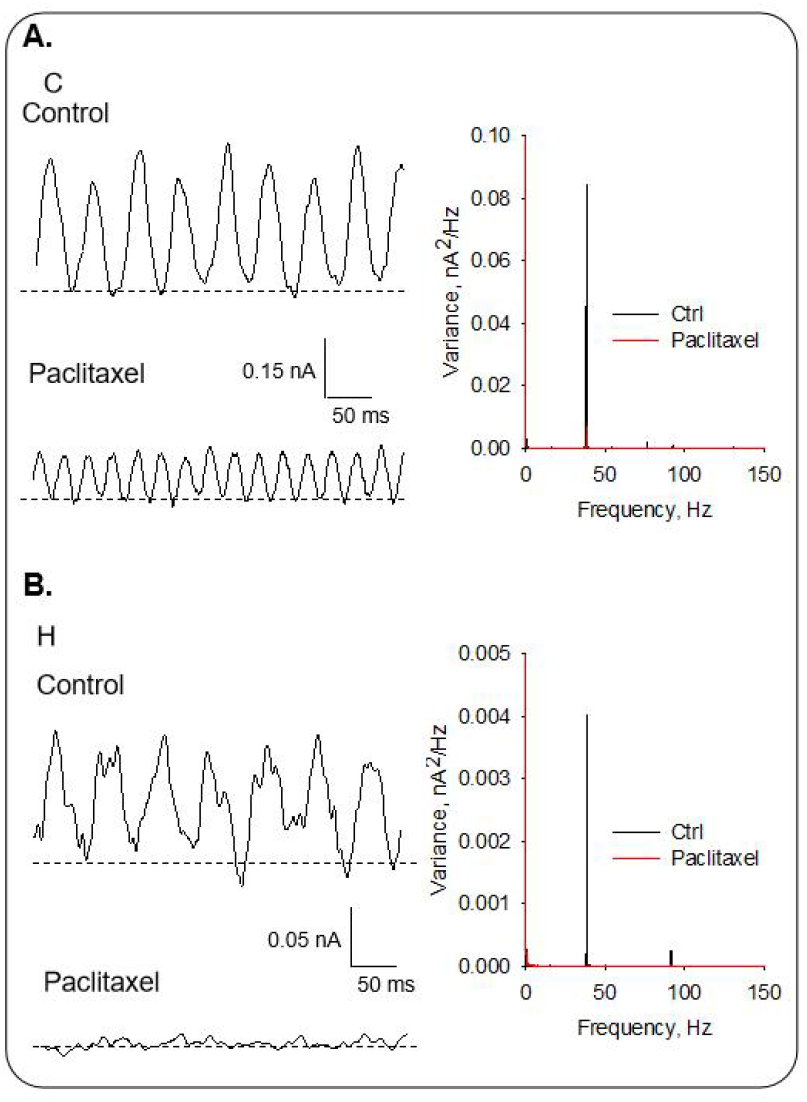
Effect of paclitaxel on rat cortex and hippocampus oscillations. a. Left. The addition of paclitaxel (1.5 mM) reduced the oscillation in cortex samples, as is shown in the tracing. Right. Fourier spectra before (Black) and after (Red) paclitaxel addition to the cortex sample. B. addition of paclitaxel inhibited almost complete oscillation in hippocampus samples. Right. Fourier spectra before (Black) and after (Red) paclitaxel addition to the hippocampus sample.

### Empirical Mode Decomposition of rat brain samples

To explore the intrinsic differences between the rat neocortex and hippocampus-generated electrical oscillations, we subjected representative electrical signals to EMD analysis, as recently reported^28^. Neocortex samples showed 5 or 6 IMFs (NaCl n = 8, KCl n = 3), while the hippocampus displayed (NaCl n = 6, KCl n = 6) 6 to 8 IMFs. Fig. 6 depicts an example of each sample in NaCl or KCl solution.

**Figure 6.**
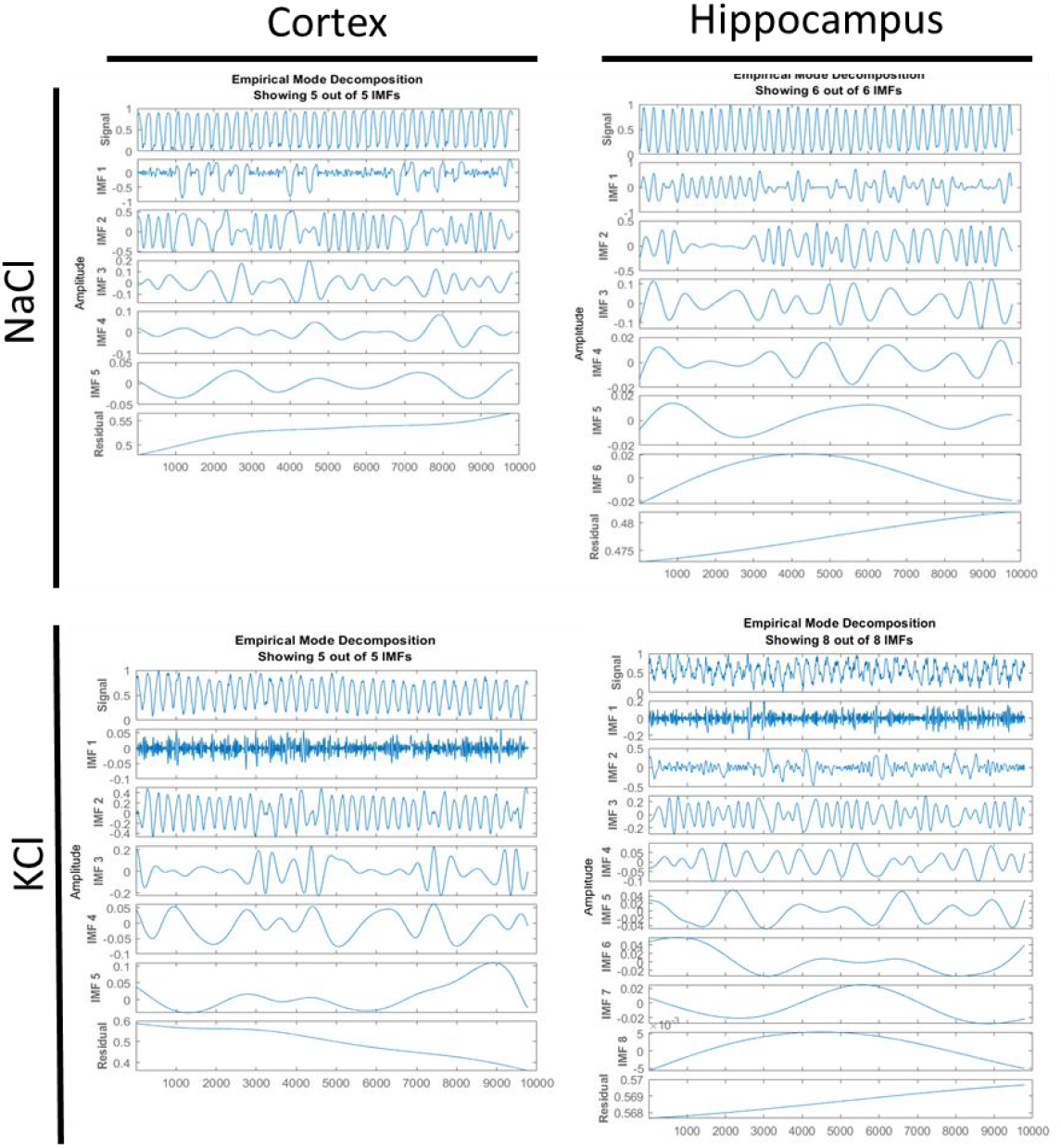
EMD of cortex and hippocampal oscillations. **(Top)** Representative EMD tracings of samples incubated in NaCl solution, for cortex (*n =* 8) and hippocampus (*n =* 6). **(Bottom)** IMFs for representative decomposition for samples incubated in KCl solution cortex (*n =* 3) and hippocampus (*n =* 6).

Each IMF obtained after the EMD was fitted to a sin function with the “Curve Fitting Tool” in Matlab 2019a. Frequencies in Hz were calculated from the Matlab parameters. Further, this strategy allowed for identifying other frequencies beyond the fundamental one.

The energy implied in each IMF was graphed using the HHT for each signal (Fig. 7A). The CWT decomposition method displays higher frequencies not seen in the HHT plots (see Fig. S1, Supplemental Material). Then, the total energy per frequency for every experiment was calculated by combining the energy implicated in each IMF (Fig. 7B), evidencing the different energy distribution in both brain areas and ion conditions.

**Figure 7.**
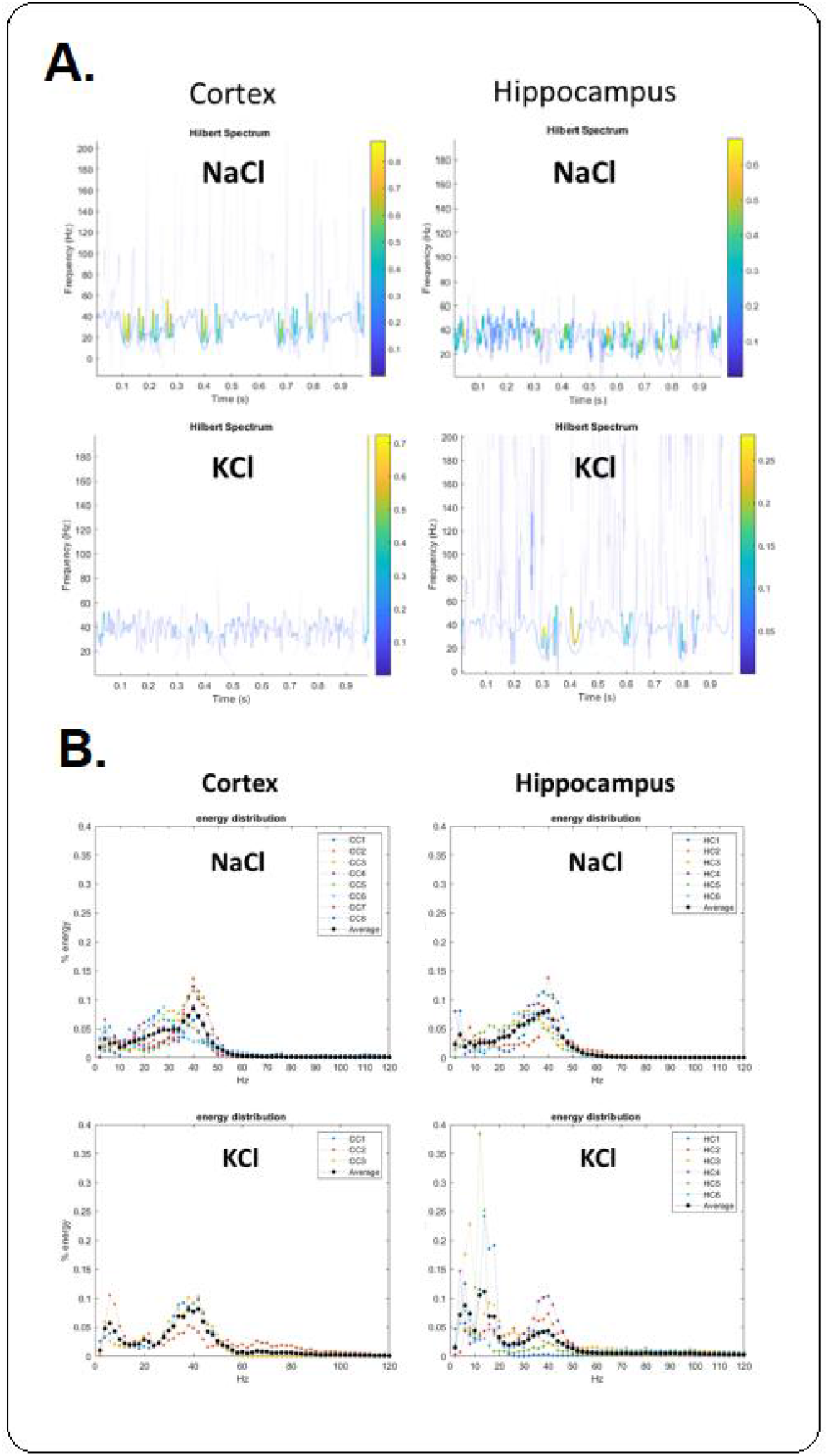
Hilbert-Huang Transform analysis of cortex and hippocampus and energy density distribution of oscillatory frequencies from cortex and hippocampus. **A**. Cortex and hippocampus HHT as indicated in the presence of either NaCl (Top) or KCl (Bottom). The figures show an example for each brain area and ion condition. **B**. Energy distribution values from individual experiments are shown in different colors, and averages are shown in black circles in NaCl (Top, cortex *n* = 8, hippocampus *n* = 6) and KCl (Bottom, cortex *n* = 3, hippocampus *n* = 6).

The energy implied in each IMF was graphed using the HHT (Fig. 7A). The CWT decomposition method displays higher frequencies not seen in the HHT plots (see Fig. S1, Supplemental Material).

### Electron microscopy of brain tissue samples

To explore the morphological correlates to the electrical properties of the brain tissue, histological samples were obtained from either the cortex or hippocampus from the same samples recorded and analyzed by electron microscopy. Samples were negatively stained, and images were obtained at 10 kV (Fig. 8). The inter-MT distances observed in the longitudinal tracks of MTs, although slightly different did not reach statistical significance (Cortex 62.08 ± 3.29 nm, n = 12, vs. 56.0 ± 2.79 nm, n = 21, p = 0.18), for the comparing cortex and hippocampus, respectively. It is important to note that in the cortex longitudinal MTs seems to be less organized than in the hippocampus. The electrical differences may be in agreement with the degree of ordering of the MT assemblies, as suggested by other MT preparations^33^. Further evidence of functional differences in the brain MTs of different locations is presented in the accompanying manuscript^31^.

**Figure 8.**
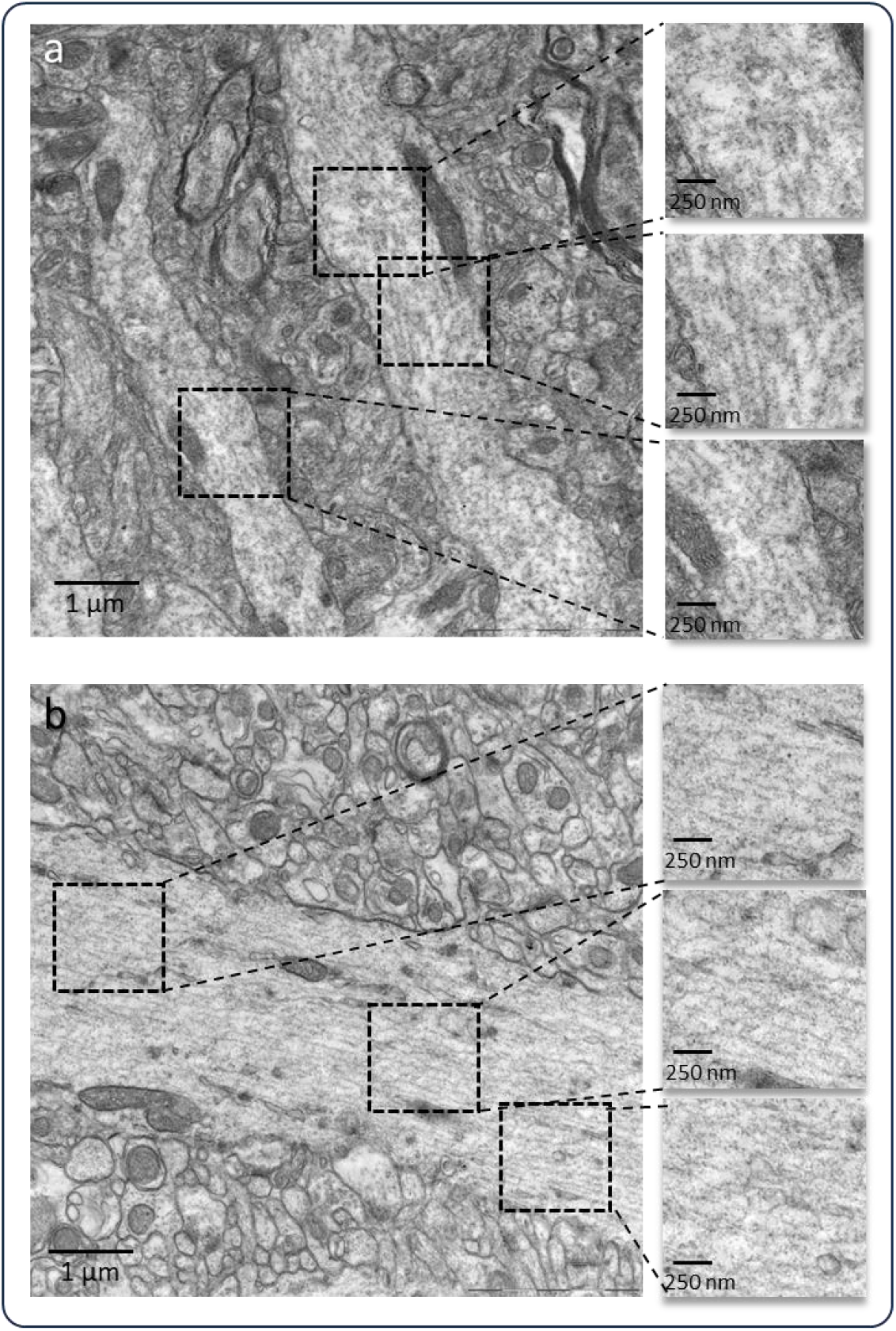
Electron microscopy of cortex (a) and hippocampus (b). Insets are shown on the Right, evidencing the MT distribution of both samples.

## Discussion

In this study, we obtained information on the electrical role of the cytoskeleton in tissue matter from the adult rat brain by applying a voltage-clamp patch pipette to assess local field potentials. The current-to-voltage relationship was approximately linear in samples incubated in the extracellular solution, and the mean conductance was statistically higher for the cortex than the hippocampus. Similar findings were obtained under KCl incubation conditions. Voltage-clamped cortex samples displayed spontaneous, self-sustained electrical oscillations that responded directly to the magnitude and polarity of the electrical stimulus. These oscillations were more prominent in the cortex samples than in those from the hippocampus. Fourier spectra showed a fundamental frequency of ∼38 Hz for both samples (cortex and hippocampus). The oscillatory frequency was present even at 0 mV, indicating that spontaneous fluctuations in the chemical gradient between the surrounding media of the brain matter and the inside of the pipette patch are sufficient to elicit electrical oscillations, as recently calculated for isolated brain MTs^32^.

Our study significantly advances understanding of the role of microtubules (MTs) in electrical oscillations. We demonstrate this intrinsic role by evaluating the effect of the MT stabilizer paclitaxel. Paclitaxel is an anti-mitotic drug, known to diffuse through, and interact with, the nanopores to binding sites in the lumen of intact MTs^34,35^. The drug’s significant suppression of the electrical oscillations in the cortex and hippocampus, consistent with its effect on other brain MT preparations, underscores the crucial role of MTs in these oscillations. Notably, the preservation of the 38 Hz frequency after paclitaxel addition further supports this conclusion.

To explore the intrinsic differences between rat neocortex and hippocampus-generated electrical oscillations, we applied Empirical Mode Decomposition (EMD) to the signals from the samples, as recently reported^28^. The neocortex samples displayed 5 or 6 Intrinsic Mode Functions (IMFs), whilst the hippocampus displayed 6 to 8 IMFs. We further evaluated the energy contributed by each IMF, using the Hilbert-Huang Transform (HHT) for each signal and the Continuous Wavelet Transform (CWT) decomposition method. These methods showed distinct properties of the waves for each brain region, tentatively correlated to the electron microscopy morphological differences in the MT order and in inter-MT distances observed in the longitudinal tracks of MTs (see Results). These differences correlated with the electrical properties of the brain tissue, suggesting a correlation with the ordering of the MT assemblies, as indicated by our findings on other MT preparations^33^. The accompanying manuscript presents further evidence of a morphological correlation with functional differences in the brain MTs of different locations^31^.

As neurons communicate, they generate electrical oscillations or brain waves, which can be measured by electroencephalography (EEG). These brain oscillations have been observed in various species, including mammals, insects, and other vertebrates and invertebrates, suggesting their fundamental role in brain computations^5,36^. There are different types of brain waves characterized by their frequency and amplitude, such as alpha, beta, delta, theta, and gamma waves^37^. Different brain regions exhibit different patterns of oscillatory activity depending on the task, or cognitive state^38^, and can communicate with each other. This, wave coherence, which is quantified as the degree of coordination between the electrical activity of different brain regions seems to be essential for various cognitive processes, including attention, memory, and perception, as it reflects the brain’s ability to integrate information across different areas. When different areas of the brain exhibit synchronized activity, they can effectively coordinate their functions, leading to enhanced cognitive performance^13,39^.

As an example of this coordination, the cortex and hippocampus are two critical brain regions that play a central role in cognitive functions. The hippocampus is primarily involved in the encoding and consolidation of new memories, whilst the cortex is responsible for the storage of long-term memories and the integration of information from various sensory inputs^40,41^. Furthermore, the coupling of patterns between hippocampal and cortical activity has been shown to facilitate memory consolidation processes. The maintenance of short-term memory involves coordinated activity between the prefrontal cortex and hippocampus, with distinct neuronal populations encoding different aspects of the memory^42^. The interconnection between the hippocampus and cortex is vital for memory consolidation, where the former transfers information to the cortex for long-term storage. This process is facilitated by synchronized activity between these regions, particularly through the coupling of oscillatory patterns^43,44^. Studies also have shown correlations in brain wave coherence between frontal and parietal regions in the alpha and beta frequency ranges. These correlations have been associated with improved attentional performance and working memory tasks^45,46^. Increased coherence between specific brain regions has been linked to improved attentional performance, while decreased coherence has been associated with attentional deficits^47^. Thus, understanding the patterns and synchronization of electrical activity in different brain regions can provide valuable insights into several brain function^48^.

Our recent studies demonstrated that brain MTs generate spontaneous, self-sustained electrical oscillations that display frequency components in the range of brain waves^28,29^. This phenomenon was observed in the cytoskeleton of permeabilized hippocampal neurons, which also display electrical oscillations^23^. MT-driven electrical oscillations with prominent peaks around 40 Hz and 90 Hz were also observed in the ex vivo honeybee brain, raising the interesting possibility that brain MTs represent a central oscillator in the brain^24^. In the present study, we explored the presence of endogenous oscillations in different identified regions of the adult rat brain. We obtained LFPs from brain tissue of the hippocampus and neocortex and observed the presence of spontaneous electrical oscillations with prominent peaks around 39 Hz and a wider region in the 18-20 Hz range. However, EMD analysis of the brain waves disclosed a different prevalence between brain locations, suggesting that the various peaks may make distinct contributions to the waveforms, and thus were only partially similar to those observed in isolated preparations of mammalian brain MTs.

### Conclusion

The present study provides direct confirmation that intracellular MTs contribute to intrinsic electrical oscillations within the cortex and hippocampus of the adult rat brain, with behavior similar to other mammalian MT preparations^23,29,49^, and also with the honeybee brain^24^. Oscillatory LFP activity in the rat brain preparations were maintained under both standard bathing solution and “intracellular-like” conditions. Spontaneous oscillations in samples of neocortex and the hippocampus suggest a widespread brain phenomenon. However, we observed exciting differences revealed by EMD analysis between these regions. While the cortex and hippocampus exhibited electrical patterns and power spectral densities, primarily peaking at frequencies around 40 Hz and 90 Hz, each contributed differently to the energy spectra. These findings align with cow and rat brain MT sheet oscillatory behavior^23,29^ but deviate somewhat from purified brain MT preparations, notably in the prominence of a 90 Hz band not apparent in cow and rat brain recordings and a distinct 18-22 Hz region observed in honeybee brain preparations^24^. This data suggests that the cytoskeleton mediates intracellular electrical signals, which may be a central phenomenon of brain tissue.

## Methods

### Animals

For the present study we used six (n = 6) adult male Wistar rats, born and reared in the breeding colony at Instituto Ferreyra (INIMEC-CONICET-UNC, Córdoba, Argentina). Animals weighing 250–300 g were housed singly in metabolic cages with free access to a regular sodium diet (Purina Rat Chow) and water *at libitum*. The room was kept at 23ºC and had a 12 h/day dark-light cycle. All experimental procedures were approved by the Ferreyra Institute’s appropriate animal care and use committee under protocol # 016/2021, following the international Public Health Service Guide for the Care and Use of Laboratory Animals (NIH Publications No. 8023, revised 1978) guidelines. We complied with the ARRIVE guidelines.

### Brain areas dissection

Immediately after decapitation, the brains were collected and placed in an ice bed for dissection. Serial coronal sections of 300 μm from the cortex (C) (bregma - 0.80 to -1.80 mm) and hippocampus (H) (bregma: -2.12 to -4.16 mm, Paxinos, and Watson, 2007) were obtained from the brain using in a vibratome. The cortex region was taken using stainless-steel needle punches of 2 mm. The extracted area included mainly the cingular cortex (Cg1 and Cg2) and part of primary and secondary motor ex (M1 and M2, respectively). The hippocampus area was separated from the cortex and slice, maintaining the hippocampus shape in each coronal section. Brain samples were washed and kept in an extracellular solution containing, in mM: 135 NaCl, 0.5 KCl, 1 MgSO_4_, 1.5 CaCl_2_, 1 EGTA, 10 HEPES, and pH adjusted to 7.23. Sections were frozen in this solution at −20°C until the time of the experiment. Note that fresh samples that were only refrigerated until the time of the experiment gave similar results to the frozen samples, indicating that freezing did not affect electrical activity. Thus, all results presented are derived from frozen samples.

### Electrophysiology

Electrical recordings from intact, ex vivo rat brains were obtained for the loose-patch-clamp configuration, as recently reported for the brain of Apis mellifera^24^. Briefly, command voltages (V_cmd_) were applied inside the brain matter of the exposed tissue. Electrode setup was similar to patch clamping with pipettes filled with a solution containing, in mM: KCl 140, NaCl 5, EGTA 1.0, and HEPES 10, adjusted to pH 7.18 with KOH. Approximately 2 mm size tissue samples were added to the dry surface of the patch clamp chamber, letting it rest for 5 min before adding 400 μl of saline solution. Experiments were conducted under symmetrical conditions, with an “external” bathing KCl solution containing the same as the pipettes^29^. Electrical recordings were conducted with a miniaturized patch-clamp amplifier, ePatch, from Elements (Cesena, Italy) with a recording range between ±200 nA, voltage stimulus range ±500 mV, and a maximum signal bandwidth of 100 kHz (Fig. 1B). Patch pipettes were made from soda lime 1.25 mm internal diameter capillaries (Biocap, Buenos Aires, Argentina) with tip diameter of ∼4 μm and tip resistance in the order of 5-15 MΩ. Voltage clamp protocols only included step-wise holding potentials (gap-free protocol) from zero mV. Electrical signals were acquired at 10 kHz and stored in a computer with the EZ patch software 1.2.11 (Elements, Cesena, Italy). Sigmaplot Version 11.0 (Jandel Scientific, Corte Madera, CA, USA) was used for statistical analysis and graphics. Power spectra of unfiltered data were obtained by the Fourier transform subroutine of Clampfit 10.0.

### Transmission electron microscopy

Samples of cortex or the hippocampus were processed following the method described previously^50^ with slight modifications for brain tissue. Briefly, tissue small sections (10-20 mm^3^) were fixed with Karnovsky’s fixative (a mixture of 2.66% w/v paraformaldehyde and 1.66% w/v glutaraldehyde in 0.1 M phosphate buffer, pH 7.2), overnight at 4°C. After fixation, cells were washed with 0.1 M phosphate buffer and embedded in 1.2% agar to form easy-to-handle sample blocks. The samples were post-fixed in 1% with osmium tetroxide for 2 hours at 4°C. After washing with phosphate buffer, they were block stained with 2% uranyl acetate for 30 minutes at room temperature in the dark. Subsequently, the samples were dehydrated in a graded ethanol series (50%, 70%, 90%, and 100%), followed by 100% acetone. Samples were then infiltrated and embedded in an acetone-SPURR resin sequence (SPURR resin; Ted Pella, Inc.). They were transferred to embedding plates and supplemented with fresh embedding medium. Polymerization was carried out at 60°C for 24 hours. After polymerization, thin sections were cut with and Powertome XL Ultramicrotome (Boeckeler Instruments, Inc.) and mounted on uncoated 200-mesh copper grids (Ted Pella). All grids were examined with a transmission electron microscope (Zeiss LIBRA 120; Carl Zeiss AG, Germany) at 80 kV, at the Electron Microscopy Core Facility and Research Center (Centro Integral de Microscopía Electrónica-CIME-CONICET-UNT).

### Drugs and chemicals

Unless otherwise stated, all reagents were obtained from Sigma-Aldrich (St. Louis, MO, USA). Wherever indicated, the MT stabilizer paclitaxel (Taxol Equivalent, Invitrogen™, P3456) was prepared as per the manufacturer’s recommendations (10 mM in DMSO) and added at the indicated concentrations.

### Empirical Mode Decomposition (EMD) Analysis

As previously reported, the EMD method was performed^28^ to analyze the data in the Time-Frequency (TF) domain. The original signal was decomposed into intrinsic mode functions (IMFs), waveform functions with only one frequency. The signals were filtered by a Gaussian filter at 200 Hz, Notch for electrical noise (50 Hz), and its harmonics using Clampfit 10.7 (Molecular Devices). Preconditioned data was parametrized using a custom function of Matlab 2019a (Mathworks, Natick, Massachusetts), and then the Matlab “emd” function was performed to obtain the signals IMFs. Each IMF coefficient (and thus its frequency) was obtained with the Matlab “Curve Fitting Tool.” The Hilbert-Huang Transform (HHT) graphics show the EMD, followed by the Huang transformation^51^ to present the IMFs with an energy-frequency-time distribution. Graphics were obtained using the Matlab function “htt”.

### Energy Calculations

Continuous Wavelet Transform (CWT) analysis is a further TF analysis that decomposes the original signal (“mother wavelet”), and the energy implied on each daughter wavelet can be calculated from the magnitude of their coefficients. Herein, we used the “cwt” Matlab function, which uses the analytic Morse wavelet with standard parameters. The CWT frequency-time plot helper was used for the graphics. Relative energy was estimated using the Fourier spectrum, calculating the area under the curve (AUC) within the range, as reported for electroencephalographic (EEG) records^52,53^. The original Fourier spectrum had a maximum frequency of 5000 Hz in all cases. A reduced frequency range was then selected to provide a Reduced Area Under the Curve (RAUC), from 0 to 140 Hz, that resembles the use of Spectral Edge Frequency (SEF) in the analysis of EEG^53^. The frequency ranges were the same as reported^28^ for comparison. The percentage of energy involved in each IMF per frequency peak was also calculated and graphed using each IMF instant frequency and energy with a 2D histogram Matlab custom function.

### Statistical analyses

Mean IMF frequency was expressed as mean and standard deviation (SD). The % RAUC was represented with mean value and standard error (SE). % RAUC per range was compared between samples using t-test or Mann-Whitney if variable normality (Shapiro Wilk) or homoscedasticity (Levenne test) failed. For statistics and graphics, SigmaPlot 11.0 software was used. Inter MT distances from TEM images were analyzed as a quantitative variable, observing mean plus minus standard error (t test), for n measurements of two different photographs. Statistical significance was set at p < 0.05.

### Schematics

The scheme shown in Fig. 1 was produced with the free software Inkscape 0.92.4 (https://www.inkscape.org).

## Acknowledgements

MdRC and HC wish to acknowledge partial funding of the present study to the Ministerio de Ciencia, Técnica, e Innovación, Argentina (PICT 0050, 2021), and CONICET, PIBAA (0495). DM thanks the Medical Research Council for generous support (MR/W028999/1).

## Author contributions statement

The authors contributions were as follows, AG, MdRC, and HFC conceived the experiments. CP and AG conducted all animal experimentation and preparation of the samples. MdRC conducted the electrophysiology experiments, curated the data, and analyzed the results. DM and ASM further analysed the results, and VHA conducted the electron microscopy studies, and HFC, MdRC, AG, ASM, and DM, designed the experiments, and prepared the draft. All authors read the manuscript.

## Additional information

Competing interests. The authors declare no competing interests

## Supplementary Material

**Table 1.** Percent RAUC for cortex and hippocampus in Na^+^ and K^+^. * *p* < 0.05 vs NaCl condition, t-test, or Mann-Whitney, as appropriate.

|  | NaCl |  |  |  |  |  | KCl |  |  |  |  |  |
| --- | --- | --- | --- | --- | --- | --- | --- | --- | --- | --- | --- | --- |
|  | Cortex |  |  | Hippocampus |  |  | Cortex |  |  | Hippocampus |  |  |
| Range | %RAUC | SE | n | %RAUC | SE | n | %RAUC | SE | n | %RAUC | SE | n |
| <2 Hz | 13.73 | 0.66 | 8 | 14.54 | 1.41 | 6 | 15.95 | 0.88 | 3 | 18.63 | 1.23 | 6 |
| 2-7 Hz | 2.86 | 0.65 | 8 | 2.99 | 0.71 | 6 | 4.72 | 1.25 | 3 | 7.36* | 1.23 | 6 |
| 9-12 Hz | 1.53 | 0.15 | 8 | 1.75 | 0.56 | 6 | 1.26 | 0.37 | 3 | 5.33* | 2.09 | 6 |
| 16-19 Hz | 2.20 | 0.29 | 8 | 2.11 | 0.61 | 6 | 2.02 | 0.66 | 3 | 2.74 | 0.37 | 6 |
| 32-44 Hz | 35.83 | 3.13 | 8 | 37.62 | 3.39 | 6 | 31.37 | 8.88 | 3 | 15.13* | 3.61 | 6 |
| 89-96 Hz | 3.17 | 0.37 | 8 | 5.07 | 1.19 | 6 | 3.44 | 0.36 | 3 | 4.40 | 0.71 | 6 |

**Fig. S1.**
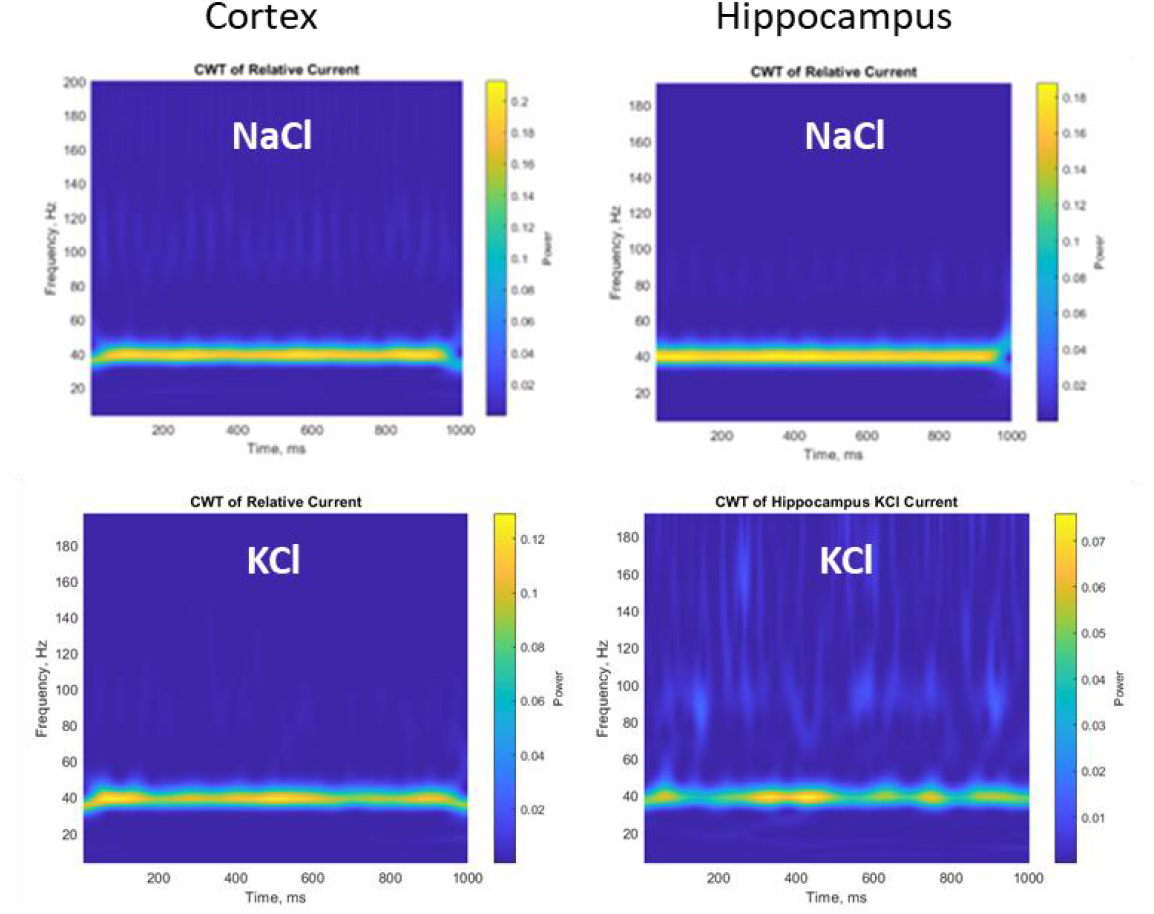
Continuous Wavelet Transform analysis of cortex and hippocampus. **Top**. Cortex (left) and hippocampus (right) CWT in NaCl. **Bottom**. Cortex (left) and hippocampus (right) CWT in KCl.

